# A High-Quality Genome Assembly from a Single, Field-collected Spotted Lanternfly (*Lycorma delicatula*) using the PacBio Sequel II System

**DOI:** 10.1101/627679

**Authors:** Sarah B. Kingan, Julie Urban, Christine C. Lambert, Primo Baybayan, Anna K. Childers, Brad S. Coates, Brian Scheffler, Kevin Hackett, Jonas Korlach, Scott M. Geib

**Affiliations:** Pacific Biosciences, Menlo Park, CA 94025, USA; Department of Entomology, The Pennsylvania State University, University Park, PA 16802, USA; USDA-ARS, Bee Research Laboratory, Beltsville, MD, 20705, USA; USDA-ARS, Corn Insects and Crop Genetics Research Unit, Ames, IA 50011, USA; USDA-ARS, Genomics and Bioinformatics Research, Stoneville, MS,38776, USA; USDA-ARS, Office of National Programs, George Washington Carver Center, Beltsville, MD, 20705, USA; USDA-ARS, Daniel K Inouye U.S. Pacific Basin Agricultural Research Center, Hilo, HI 96720, USA

**Author notes:** Correspondence address. Scott Geib, USDA-ARS, Daniel K Inouye U.S. Pacific Basin Agricultural Research Center, 64 Nowelo St., Hilo, HI 96720; Jonas Korlach, Pacific Biosciences, 1305 O’Brien Drive, Menlo Park, CA 94025.

## Abstract

A high-quality reference genome is an essential tool for applied and basic research on arthropods. Long-read sequencing technologies may be used to generate more complete and contiguous genome assemblies than alternate technologies, however, long-read methods have historically had greater input DNA requirements and higher costs than next generation sequencing, which are barriers to their use on many samples. Here, we present a 2.3 Gb *de novo* genome assembly of a field-collected adult female Spotted Lanternfly (*Lycorma delicatula*) using a single PacBio SMRT Cell. The Spotted Lanternfly is an invasive species recently discovered in the northeastern United States, threatening to damage economically important crop plants in the region. The DNA from one individual was used to make one standard, size-selected library with an average DNA fragment size of ~20 kb. The library was run on one Sequel II SMRT Cell 8M, generating a total of 132 Gb of long-read sequences, of which 82 Gb were from unique library molecules, representing approximately 36-fold coverage of the genome. The assembly had high contiguity (contig N50 length = 1.5 Mb), completeness, and sequence level accuracy as estimated by conserved gene set analysis (96.8% of conserved genes both complete and without frame shift errors). Further, it was possible to segregate more than half of the diploid genome into the two separate haplotypes. The assembly also recovered two microbial symbiont genomes known to be associated with *L. delicatula*, each microbial genome being assembled into a single contig. We demonstrate that field-collected arthropods can be used for the rapid generation of high-quality genome assemblies, an attractive approach for projects on emerging invasive species, disease vectors, or conservation efforts of endangered species.

## Background

In September 2014, *Lycorma delicatula* (Hemiptera: Fulgoridae), commonly referred to as the Spotted Lanternfly, was first detected in the United States in Berks County, Pennsylvania. *L. delicatula* is a highly polyphagus phloem-feeding insect native to Asia that is documented to feed upon more than 65 plant species [1, 2]. Because this insect was an invasive that damaged grape vines and tree fruit in South Korea in the mid-2000s [3, 4], its potential to cause economic damage was known. Shortly after it was detected in the U.S., the Pennsylvania Department of Agriculture established a quarantine zone surrounding the site of first detection. The invasion likely began with a shipment of stone that harbored egg masses, as *L. delicatula* lays inconspicuous egg masses seemingly indiscriminately on a wide variety of surfaces (e.g., tree bark, automobiles, rail cars, shipping pallets, etc.), contributing to the potential for abrupt and distant spread. Since that time, the *L. delicatula* quarantine zone has expanded from an area of 50 mi^2^ to over 9,400 mi^2^. While this pest has huge potential for spread and increased impact, essentially nothing is known at the genomic level about this species or any Fulgorid species, and there is a need to develop resources rapidly for this pest to support development of management and control practices.

A high-quality genome as a foundation to understand arthropod biology can be a powerful tool to combat invasions and disease-carrying vectors, aid in conservation, and many other fields (for examples, see [5–8]). To this end, large-scale initiatives are underway to comprehensively catalog the genomes of many arthropod species, including the i5K initiative aiming to sequence and analyze the genomes of 5,000 arthropod species [9–11] associated with the Darwin Tree of Life Project (https://www.sanger.ac.uk/news/view/genetic-code-66000-uk-species-be-sequenced), and the Earth BioGenome Project [12]. Within the context of the Earth BioGenome Project, the USDA-ARS Ag100Pest initiative is focused on rapidly deciphering the genomes of 100 destructive insect species to crops and livestock, projected to have profound bioeconomic impacts to agriculture and livestock industries, as well as habitat and species conservation.

Arthropod genome assembly projects face unique challenges stemming from their small body size and high heterozygosity. Due to the limited quantities of genomic DNA that can be extracted from a small-bodied animal, researchers may pool multiple individuals, such as by generating NGS libraries of different insert sizes, each from a different individual [10, 13], or by pooling multiple individuals for a single long-read sequencing library from an iso-female laboratory strain [14–18] or colony [5, 19]. Pooling introduces multiple haplotypes into the sample and complicates the assembly and curation process [19], and while this issue may be ameliorated by inbreeding it is not always an option for organisms that cannot be cultured in the laboratory. Moreover, genomic regions with high heterozygosity tend to be assembled into more fragmented contigs [20], so computational methods specifically developed for heterozygous samples are needed [21–23]. Recently, high-quality long-read assemblies have been published for a single diploid mosquito (*Anopheles coluzzii*) [24] and a single haploid honeybee (*Apis mellifera*) [7]. Despite both species having relatively small genomes (<300 Mb), multiple PacBio SMRT Cells were needed for sufficient sequencing coverage (N=3 for mosquito, N=29 for bee).

Here, we demonstrate the sequencing and high-quality *de novo* assembly of a 2.25 Gb genome from a single, field-collected Spotted Lanternfly (*Lycorma delicatula*) insect, requiring only one sequencing library and one SMRT Cell sequencing run on the Sequel II System. The genome assembly is highly contiguous, complete and accurate, and resolves the maternal and paternal haplotypes over 60% of the genome. In addition to the lanternfly genome, the assembly immediately provided complete genomes from two of the organism’s bacterial endosymbionts. The approach outlined here can be applied to field-collected arthropods or other taxa for which the rapid generation of high-quality contig-level genome assemblies is critical, such as for invasive species or for conservation efforts of endangered species.

## Results

We extracted DNA from a single female *L. delicatula* collected from the main trunk of *Ailanthus altissima* (tree of heaven) in Reading, Berks County, Pennsylvania, USA (40.33648 N, 75.90471 W) on the 26th of August 2018 (Figure 1). *L. delicatula* is known to harbor several endosymbionts in specialized bacteriocytes, predominantly in the distal end of the insect abdomen; to avoid a high proportion of these symbionts in the sequencing, DNA was extracted from the head and thorax regions of the insect only (see Materials & Methods for details). While more recently developed single arthropod assemblies have significantly lowered DNA input requirements [24], here the amount of extracted genomic DNA was more plentiful because of the relatively larger size, allowing for sufficient DNA for a standard library preparation with size selection, resulting in a ~20 kb average insert size sequencing library (Figure 2). The library was sequenced on the Sequel II System with one SMRT Cell 8M, yielding 131.6 Gb of total sequence contained in 5,639,857 reads, with a polymerase read length N50 of 41.7 kb and insert (subread) length N50 of 22.3 kb (Figure S1).

**Figure 1.**
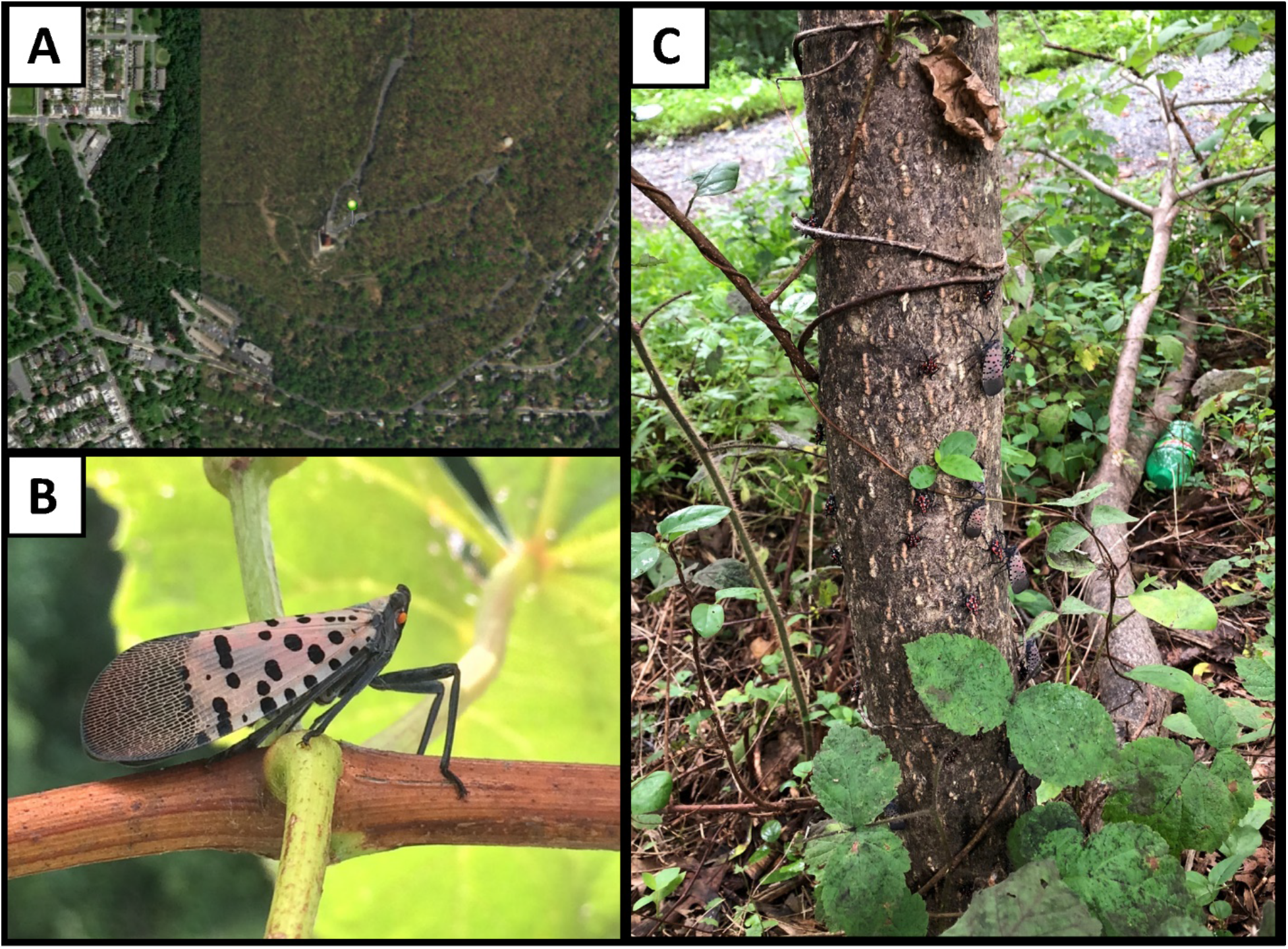
Specimen collection. (A) Location of specimen collection (green marker), near the Reading Pagoda on Mt. Penn (Reading, Berks County, Pennsylvania, USA (40.33648 N, 75.90471 W)); (B) Adult female *Lycorma delicatula*; (C) The host *Ailanthus altissima* tree (tree of heaven) from which the adult female sample was collected on 26th of August 2018. Late nymph stage and adults can be seen covering the trunk of this host tree.

**Figure 2.**
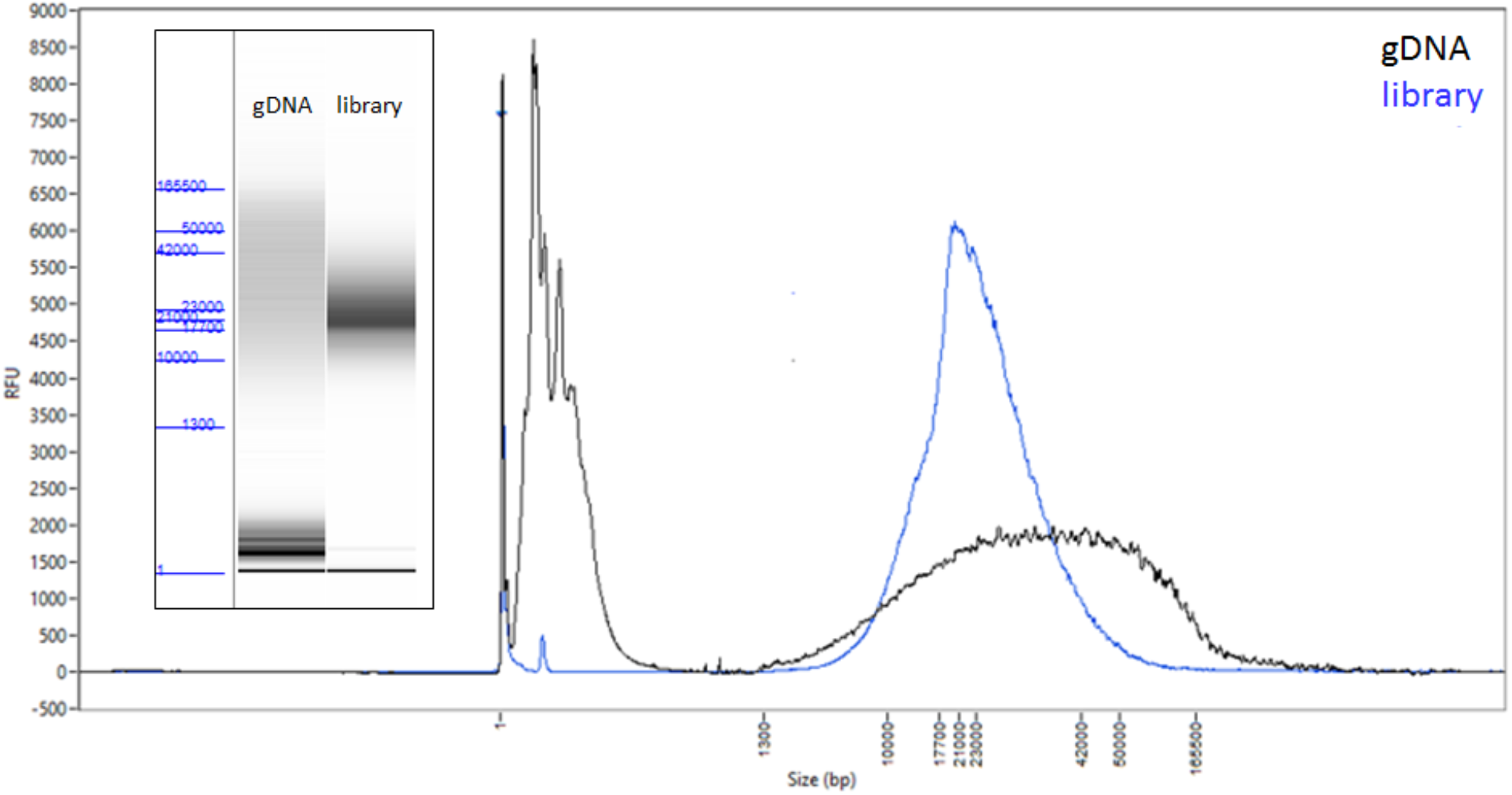
*Lycorma delicatula* input DNA and resulting library. FEMTO Pulse traces and ‘gel’ images (inset) of the genomic DNA input (black) and the final library (blue) before sequencing.

The genome was assembled with FALCON-Unzip, a diploid assembler that captures haplotype variation in the sample [21]. A single subread per ZMW was used in assembly for a total of 82.4 Gb of sequence (36-fold coverage for a 2.3 Gb genome). Reads longer than 8 kb were selected as “seed reads” for pre-assembly, a process of error correction using alignment and consensus calling with the PacBio data. Pre-assembled reads totaled 55.5 Gb of sequence (24-fold) with mean (N50) read length of 10.8 kb (15.2 kb) (Figure S1). The draft FALCON assembly consisted of 5,158 contigs with N50 length of 1.38 Mb and total assembly size of 2.43 Gb. We screened this draft assembly for bacterial symbiont or contaminant DNA (see methods) and identified two contigs originating from microbial symbionts, *Sulcia muelleri* and *Vidania fulgoroideae*, respectively, two known bacterial symbionts of planthoppers [25]. These contigs were removed from the final curated assembly and analyzed separately (see below).

The FALCON-Unzip module was applied to phase and haplotype-resolve the assembly. The unzipped assembly was then polished twice to increase base-level accuracy of the contigs, using all subreads, including multiple passes from a single library molecule. The resulting assembly consisted of 4,209 primary contigs comprising 2.40 Gb with contig N50 of 1.42 Mb. A total of 1.25 Gb of the assembly “unzipped” into 10,103 haplotigs of mean (N50) length 76.9 kb (152 kb) (Table 1).

**Table 1:** Spotted Lanternfly *de novo* genome assembly stats for the FALCON-Unzip and curated assemblies. Assembly contiguity and BUSCO completeness stats are shown after FALCON-Unzip, and after curation to recategorize duplicated haplotypes in the primary contigs, removal of repetitive and redundant haplotigs and bacterial contigs. For complete BUSCO stats see Table S1.

| Assembly Version | FALCON-Unzip | Curated Assembly |
| --- | --- | --- |
| Primary assembly size | 2.395 Gb | 2.252 Gb |
| Number of primary contigs | 4209 | 2927 |
| Contig length N50 | 1.423 Mb | 1.520 Mb |
| Haplotig assembly size<br>(proportion of primary length) | 1.249 Gb (52%) | 1.349 Gb (60%) |
| Number of haplotigs | 10,103 | 10,652 |
| Haplotig N50 | 185.5 kb | 178.1 kb |
| BUSCO complete | 96.7 % | 96.7 % |
| BUSCO duplicate | 3.3 % | 2.4 % |

While FALCON-Unzip is designed to resolve haplotypes in non-inbred organisms, some homologous regions of the genome with high heterozygosity may be assembled on separate primary contigs. Our goal was to generate a haploid reference sequence, so we performed additional curation to both recategorize duplicated haplotypes from the primary set as haplotigs and remove repetitive, artefactual, and redundant haplotigs (see Methods). The final curated assembly consisted of 2,927 primary contigs of total length 2.25 Gb with contig N50 1.52 Mb. The alternate haplotypes spanned 60% of the primary contig length: 10,652 haplotigs comprised a total of 1.35 Gb with an N50 length of 178 kb. A visualization of the assembly contiguity and completeness was generated using assembly-stats [26] and are presented in Figure 3 and Table 1.

**Figure 3.**
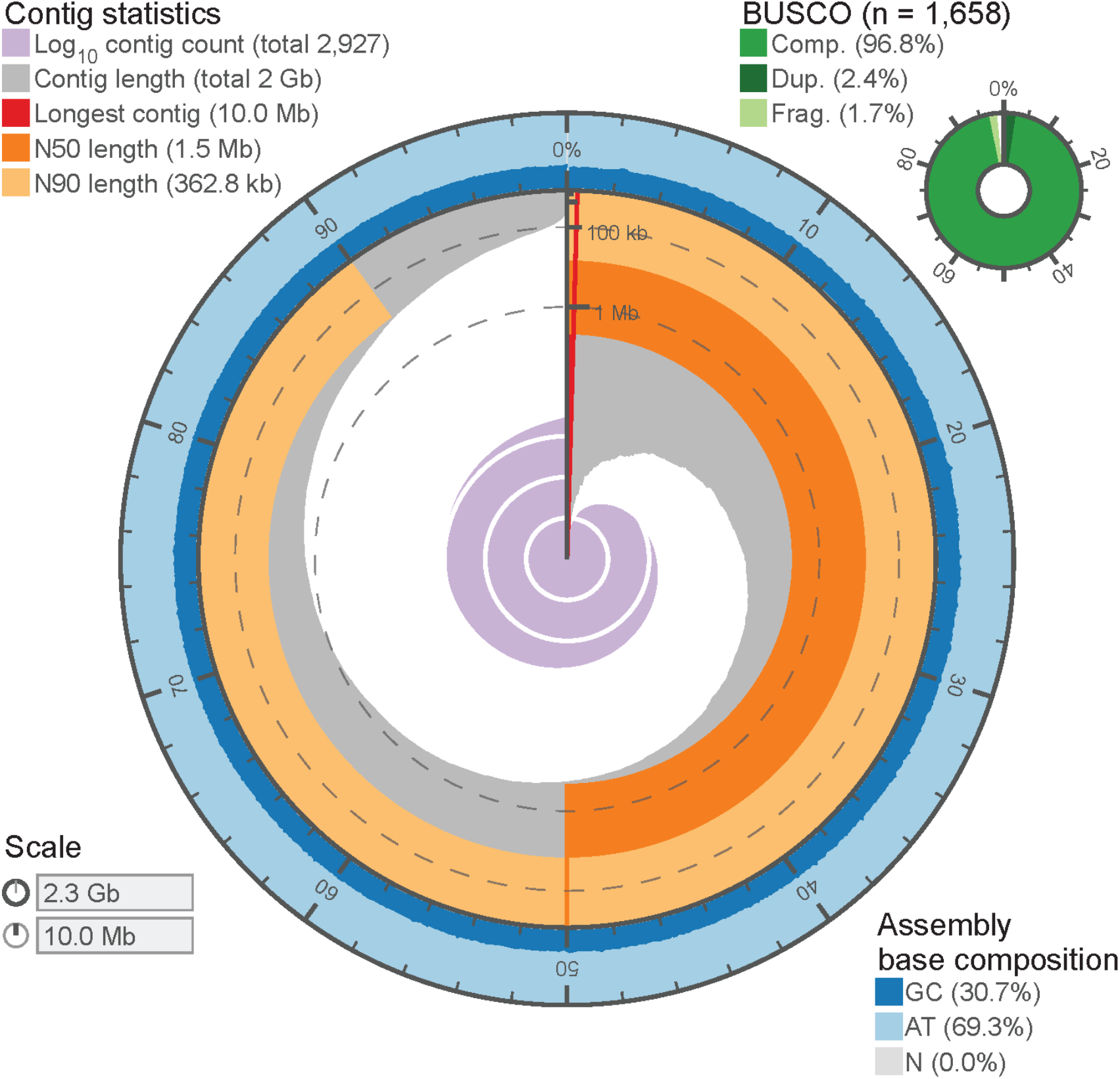
Assembly Visualization. The contiguity and completeness of the *L. delicatula* genome assembly is visualized as a circle, with the full circle representing the full assembly length of ~2.3 Gb. The longest contig was 10.0 Mb, and the assembly has uniform GC content throughout, with very few contigs below 50 kb in length.

Despite attempting to avoid bacteriocyte associated internal symbionts by excluding the abdomen during DNA extraction, two contigs which were identified as circular and of microbial origin were present in the assembly. Contig 001940F is a complete representation of the Candidatus *Sulcia muelleri* obligate symbiont, 212,195 bp in length with a GC content of 23.8% and sequenced at approximately 46.6-fold coverage of subreads. A second contig 5193, designated to be circular in the FALCON assembly, was identified as a complete representation of the Candidatus *Vidania fulgoroideae* obligate symbiont genome. This genome was 126,523 bp in length with a GC content of 19.15%. Contig names are relative to the FALCON assembly prior to running Unzip, which is available in the supporting dataset on the Ag Data Commons (see Data Availability). More details on these symbionts will be provided in an accompanying manuscript.

We assessed additional aspects of genome assembly completeness and sequence accuracy with analysis of conserved genes. Using the ‘insecta_odb9’ BUSCO gene set collection [27], we observed >96% of the 1,658 genes were complete and >96% occurred as single copies (Tables 1 & S1). Concordantly with the recategorization of initial primary contigs into haplotigs by Purge Haplotigs, the percentage of duplicated genes decreased from 3.3% to 2.4%. As an additional evaluation, we aligned to the primary assembly the core *Drosophila melanogaster* CEGMA gene set, resulting in 416 alignments (91%) and an average alignment length of 86%, and with >96.6% of alignments showing no frame shift-inducing indels.

## Discussion and Conclusions

We sequenced and assembled a high-quality reference genome for a single wild-caught Spotted Lanternfly (*Lycorma delicatula*), a Fulgorid planthopper species invasive in the northeastern U.S. Previous planthopper genome projects required 100-5,000 inbred individuals and at least 16 different sequencing libraries [28–30] (Table 2). We generated long-read sequence data sufficient for *de novo* assembly from a single sequencing library, run on one PacBio SMRT Cell. Despite the fact that the genome of our planthopper species is 2-4 times larger compared to the three previous described planthopper genomes, it is 13 to 63 times more contiguous. The new workflow presented here improves on many aspects of previous approaches for generating arthropod genome assemblies, and the genomes of their endosymbionts. These include sample (i) collection strategies, (ii) library preparation efforts and sequencing time, (iii) assembly considerations, and (iv) endosymbiont genome capture and are discussed in detail below.

**Table 2:** Comparison of the Spotted Lanternfly genome assembly with previously described planthopper species assemblies. Current study has improvements with regard to the required number of insect individuals, sequencing libraries, assembly sizes and contiguity qualities. ^a^[29]; ^b^[28] & Q.Wu personal communication; ^c^[30]& F. Cui personal communication.

| Species | *Nilaparvata lugens*<br>(2014)<sup>a</sup> | *Sogatella furcifera*<br>(2017)<sup>b</sup> | *Laodelphax striatellus*<br>(2017)<sup>c</sup> | *Lycorma delicatula*<br>(this work) |
| --- | --- | --- | --- | --- |
| Number of Individuals<br>(source) | ~5,000<br>(F13 from inbred line) | ~120<br>(F6 from inbred line) | ~100<br>(F22 from inbred line) | 1<br>(field-collected) |
| Number of Sequencing<br>Libraries | 16<br>(+fosmid libraries) | 17 | 47 | 1 |
| Assembly Size | 1.14 Gb | 0.72 Gb | 0.54 Gb | 2.25 Gb |
| Contig N50 | 24 kb | 71 kb | 118 kb | 1,520 kb |

### Collection strategies

The strategy of performing single-insect genome assemblies has several advantages. First, it dispenses with the requirement of inbred lab colonies, which may take months or even years to establish, can be expensive to maintain, and are impractical or impossible for many species. Second, by sampling field-collected animals, genetic variation can be more accurately characterized for local populations, without the risk of adaptation to lab culture [31] or loss of heterozygosity [32]. For invasive pests, methods for artificial rearing often do not exist and there is a desire to rapidly generate foundational data on these pests, so direct sequencing of wild specimens is advantageous. The ability to generate genomes *de novo* from field-collected arthropods makes high-quality genomes accessible for many more species. This approach also enables comprehensive comparisons of genetic diversity within and between populations without the bias from single reference-based studies [15] and allows generation of a diploid genome assembly that more closely captures the organism’s biology [22].

### Library preparation and sequencing

The methods described here for DNA extraction, library preparation and sequencing are straightforward and rapid, using established kits and leveraging the higher throughput of the Sequel II System to generate sufficient sequencing coverage with just one SMRT Cell and 30 hours of sequencing run time. The need for multiple libraries from several individuals or pool fractions, or for covering different insert size ranges is eliminated. These improvements potentially allow a genome project, with infrastructure optimization, to be completed in less than one week (estimating one day each for DNA extraction, library preparation, sequencing, and data analysis), and can be carried out by individual labs rather than requiring large consortia that were typical of previous genome assembly efforts. All steps in the workflow are amenable to automation to accommodate larger sample numbers in a high throughput manner. The rapid nature of the workflow will allow for not only the generation of a single reference-grade genomic resource, but for the comprehensive genomic monitoring of species before or throughout a field season, and for rapid testing of intervention strategies.

### Assembly

An additional advantage to single-insect assemblies is that genome assembly for a diploid sample is algorithmically simpler than for a sample of many pooled individuals, each of which may contribute up to two unique haplotypes. Several *de novo* assembly methods are available for diploid samples [21, 22, 33], and have been broadly applied taxonomically [5, 34]. Recent work indicates that assembly of high-heterozygosity samples is more accurate than for inbred samples when parental data can be used to partition long-read sequence data by haplotype, an approach called trio-binning [22, 35]. When trio samples are not available, long-range contact data may be leveraged in combination with long-read assemblies to enhance haplotype phasing [36]. This represents a reversal in the paradigm for high-quality references in insect genomics [22], where one now should target outcrossed or highly heterozygous (wild) individuals, rather than inbreeding to reduce polymorphism and avoid complications caused by heterozygosity that may arise using previous assembly methods.

### Endosymbionts and metagenomic approaches for symbiosis

Although our method of DNA extraction was intended to avoid structures in the lanternfly that house bacterial symbionts, our results included the complete genomes of two known planthopper endosymbionts, *Sulcia muelleri* and *Vidania fulgoroideae*. Early work by Müller revealed that the cells (bacteriocytes) housing endosymbionts in planthoppers are organized into organs, or bacteriomes, and that these structures often display complex morphologies and occupy a variety of positions within an insect’s abdomen [37]. Dissections of *L. delicatula* reveal the presence of complex, string-like bacteriome structures positioned around the alimentary canal that are large enough to be visible to the naked eye. As such, it is not surprising that some bacteriome tissue was included with the thorax as it was separated from the abdomen for extraction. Despite the attempt to avoid these symbionts, their complete genomes were recovered at sufficient coverage to be assembled into single contigs from a host-targeted DNA extraction. This approach allows for high-quality assemblies in a metagenomic context, with the long reads and robust assembly strategy allowing for clear discrimination of the microbial symbionts. This dramatically simplifies strategies for symbiont sequencing: rather than dissection and pooling of bacteriocytes from the host, a shotgun metagenomics strategy can be used to not only recover the symbiont genome but also a draft reference of the host, at a similar cost to targeted methods. Additional follow-up shotgun approaches could yield discovery of novel or unexpected microbes associated with the host.

### Genomic applications for control

High-quality reference genomes for *L. delicatula* and its associated endosymbionts represent invaluable resources for this dangerous invasive, about which little is known of its basic biology. Because obligate symbionts in phloem-feeding insects typically provide nutritional benefit to their hosts [38, 39], the symbiont genomes offer insight into nutritional requirements and basic metabolic functioning of *L. delicatula*. They also offer additional potential opportunities for control. For example, obligate endosymbionts are typically vertically transferred from female to offspring transovarially. In *L. delicatula*, development of the female reproductive system appears to require substantial time and resources. Females typically eclose as adults in late July, and feed voraciously over several months and accumulate abdominal mass, before laying eggs in October-November. During this time, bacterial symbionts must proliferate and get transferred to developing ovarioles. This may present a time window for potential disruption of symbiont transmission, which would represent a control strategy that is highly specific to *L. delicatula*. Alternatively, RNA inhibition (RNAi) strategies used for control often target highly conserved genes in the insect’s genome that perform vital cellular functions. Inhibition of one of these core gene functions is lethal to the insect. Targeting such highly conserved genes, however, reduces the species specificity of this approach. Obligate bacterial endosymbionts, however, only occur within the host insects with whom they have coevolved over tens to hundreds of millions of years, and as such, provide highly species-specific genomic targets for control with RNAi [40].

### Conclusions

The genome assembly presented here can be used as a foundation for further assembly and curation efforts with long-range scaffolding technologies such as Bionano Genomics [41, 42] and/or Hi-C [19, 43–45] to generate a reference-quality, chromosome-scale genome scaffold representation. Similarly, full-length RNA-seq (Iso-Seq) [46, 47] or other RNA-seq data types can be applied, with the assembly serving as a mapping reference, for gene and other functional element annotation. While these follow-up efforts are currently underway in our laboratories, we wanted to make this initial, high-quality draft genome assembly available in the hope that it will provide a valuable resource to the scientific community to improve our understanding for this important agricultural pest.

## Materials & Methods

### Sample collection and DNA isolation

A cohort of *L. delicatula* females were collected off of their preferred host *Ailanthus altissima* (tree of heaven) in Reading, Berks County, Pennsylvania, USA (40.33648 N, 75.90471 W) on the 26^th^ of August 2018. Individuals were snap frozen in liquid nitrogen in the field and stored at −80 °C until processing. *L. delicatula* were extracted individually, by first cutting off the abdomen, and grinding the head and thorax in liquid nitrogen to a powder. High molecular weight DNA was extracted using a modification of a “salt-out” protocol described (https://support.10xgenomics.com/de-novo-assembly/sample-prep/doc/demonstrated-protocol-dna-extraction-from-single-insects). Briefly, the ground material was resuspended in 1.8 ml of lysis buffer (10 mM Tris-HCl, 400 mM NaCl, and 100 mM EDTA, pH 8.0) and 120 μl of 10% SDS and 300 ul of Proteinase K solution (1 mg/ml Proteinase K, 1% SDS, and 4 mM EDTA, pH 8.0) was added. The sample was incubated overnight at 37 °C. To remove RNA, 40 μl of 20 mg/ml RNAse A was added and the solution was incubated at room temperature for 15 minutes. Seven hundred and twenty μl of 5 M NaCl was added and mixed gently through inversion. The sample was centrifuged at 4 °C at 1500 × g for 20 minutes. A wide-bore pipette tip was then used to transfer the supernatant, avoiding any precipitated protein material, to a new tube and DNA was precipitated through addition of 3.6 ml of 100% EtOH. The DNA was pelleted at 4 °C at 6250 × g for 15 min, and all EtOH was decanted from the tube. The DNA pellet was allowed to dry and then was resuspended in 150 μl of TE. Initial quality and quantity of DNA was determined using a Qubit fluorometer and evaluating DNA on a 1% agarose genome on a Pippin Pulse using a 14-hour 5kb - 80kb separation protocol. DNA was sent to Pacific Biosciences (Menlo Park, California) for library preparation and sequencing.

### Library preparation and sequencing

Genomic DNA quality was evaluated using the FEMTO Pulse automated pulsed-field capillary electrophoresis instrument (Agilent Technologies, Wilmington, DE), showed a DNA smear, with majority >20kb (Figure 2), appropriate for SMRTbell library construction without shearing.

One SMRTbell library was constructed using the SMRTbell Express Template Prep kit 2.0 (Pacific Biosciences, Menlo Park, CA). Briefly, 5 μg of the genomic DNA was carried into the first enzymatic reaction to remove single-stranded overhangs followed by treatment with repair enzymes to repair any damages that may be present on the DNA backbone. After DNA damage repair, ends of the double-stranded fragments were polished and subsequently tailed with an A-overhang. Ligation with T-overhang SMRTbell adapters was performed at 20 °C for 60 minutes. Following ligation, the SMRTbell library was purified with 1X AMPure PB beads. The size distribution and concentration of the library were assessed using the FEMTO Pulse and dsDNA BR reagents Assay kit (Thermo Fisher Scientific, Waltham, MA). Following library characterization, 3 μg was subjected to a size-selection step using the BluePippin system (Sage Science, Beverly, MA) to remove SMRTbells ≤ 15 kb. After size selection, the library was purified with 1X AMPure PB beads. Library size and quantity were assessed using the FEMTO Pulse (Figure 2), and the Qubit Fluorometer and Qubit dsDNA HS reagents Assay kit.

Sequencing primer v2 and Sequel II DNA Polymerase were annealed and bound, respectively, to the final SMRTbell library. The library was loaded at an on-plate concentration of 30 pM using diffusion loading. SMRT sequencing was performed on the Sequel II System with Sequel II Sequencing Kit, 1800-minute movies, and Software v6.1.

### Assembly

Data were assembled with FALCON-Unzip [21] using pb-falcon version 0.2.6 from the bioconda pb-assembly metapackage version 0.0.4 with the following configuration:

genome_size = 2500000000; seed_coverage = 30; length_cutoff = −1; length_cutoff_pr = 10000; pa_daligner_option = −e0.8 -l 1000 -k18 -h70 -w8 -s100; ovlp_daligner_option = -k24 -h1024 - e.92 -l 1000 -s100; pa_HPCdaligner_option = -v -B128 -M24; ovlp_HPCdaligner_option = -v - B128 -M24; pa_HPCTANmask_option = -k18 -h480 -w8 -e.8 -s100; pa_HPCREPmask_option = -k18 -h480 -w8 -e.8 -s100; pa_DBsplit_option = -x500 -s400; ovlp_DBsplit_option = -s400; falcon_sense_option = --output-multi --min-idt 0.70 --min-cov 3 --max-n-read 100 --n-core 4; overlap_filtering_setting = --max-diff 100 --max-cov 200 --min-cov 3 --n-core 24; polish_include_zmw_all_subreads = true

The assembly was polished once as part of the FALCON-Unzip workflow and a second time by mapping all subreads to the concatenated primary and haplotig reference with pbmm2 v1.1.0 (“pbmm2 align $REF $BAM $MOVIE.aln.bam --sort -j 48 -J 48”) and consensus calling with arrow using gcpp v 0.0.1-e2ea76a (“gcpp -j 4 -r $REF -o $OUT.$CONTIG.fasta $BAM -w “$W””). Both tools are available through bioconda: https://github.com/PacificBiosciences/pbbioconda. We screened the primary assembly for duplicate haplotypes using Purge Haplotigs (bioconda v1.0.4) [48]. Purge Haplotigs identifies candidate haplotigs in the primary contigs using PacBio read coverage depth and contig alignments. To determine the coverage thresholds, we mapped only the unique subreads to the primary contigs rather that all subreads. This resulted in more distinct modes in the coverage histogram (data not shown). A fasta file of unique subreads was generated with the command, “python -m falcon_kit.mains.fasta_filter median movie.subreads.fasta > movie.median.fasta” which is available in the pb-assembly software. We used coverage thresholds of 5, 25, and 10 and default parameters except “-s 90” (diploid coverage maximum for auto-assignment of contigs as suspect haplotigs). We recategorized 1,269 primary contigs as haplotigs (total length 141.8 Mb), discarded 12 as artifactual (total length 869 kb) and 201 as repeats (total length 19.1 Mb). A perl script (https://github.com/skingan/adapt_PurgeHaplotigs_for_FALCONPhase) was used to rename the haplotigs using the FALCON-Unzip nomenclature so that each haplotig can be easily associated with a primary contig. Following renaming, we aligned each haplotig to its associate primary contig, chained sub-alignments in one dimension, and removed redundant haplotigs whose alignment to the primary was completely contained within another haplotig [36]. This process removed 518 haplotigs totaling 22.6 Mb.

### Contaminant and symbiont screening

All primary contigs from the draft FALCON assembly were searched using DIAMOND BLASTx against the NCBI nr database (downloaded April 8th, 2019) [49], and the subsequent hits were used to assign taxonomic origin of each contig using a least common ancestor assignment for each contig utilizing MEGAN 6.15.2 Community Edition with the longReads LCA Algorithm and readCount assignment mode [50]. Any contigs that were identified as microbial were flagged and removed from the final assembly. To avoid assignment of contigs as microbial when a microbial gene may have horizontally transferred to the insect, any potentially microbial contigs were screened for presence of BUSCO insect genes and retained if a BUSCO was present on the contig.

### Genome assembly evaluation

To assess the completeness of the curated assembly, we searched for conserved, single copy genes using BUSCO (Benchmarking Universal Single-Copy Orthologs) v3.0.2 [27] with the ‘insecta_odb9’ database. In addition, we evaluated assembly completeness and accuracy against the *Drosophila melanogaster* CEGMA gene set (http://korflab.ucdavis.edu/datasets/cegma/core_genome/D.melanogaster.aa), using a previously described script [51]. A visualization of the assembly contiguity and completeness was generated using assembly-stats [26] and are presented in Figure 3 and Table 1.

## Availability of Data

Raw data and final assembly for this project are submitted to NCBI under BioProject PRJNA540533, sample described in BioSampleSAMN11546444, SRA accession for raw PacBio subreads is SRR9001434. Supporting data to this publication is submitted to the AgDataCommons at https://data.nal.usda.gov/dataset/data-high-quality-genome-assembly-single-field-collected-spotted-lanternfly-lycorma-delicatula-using-pacbio-sequel-ii-system under DOI: 10.15482/USDA.ADC/1503745, including polished FALCON assembly, polished FALCON-Unzip assembly, final curated assembly and placement file, microbial symbiont assemblies and associated metadata.

## Additional Files

**Table S1:** Full summary from BUSCO analysis of primary contigs and haplotigs. The ‘insecta_odb9’ gene set was used (Total genes = 1658), after different stages of assembly and curation.

| **Gene Count** | **FALCON-Unzip**<br>**Primary (haplotigs)** | **Purge Haplotigs**<br>**Primary (haplotigs)** | **Final Curated**<br>**Primary (haplotigs)** |
| --- | --- | --- | --- |
| **Complete** | 1602 (954) | 1604 (977) | 1604 (975) |
| **Duplicate** | 54 (12) | 40 (16) | 40 (14) |
| **Single Copy** | 1548 (942) | 1564 (961) | 1564 (961) |
| **Fragmented** | 33 (110) | 30 (99) | 30 (101) |
| **Missing** | 23 (594) | 24 (582) | 21 (582) |

**Figure S1:**
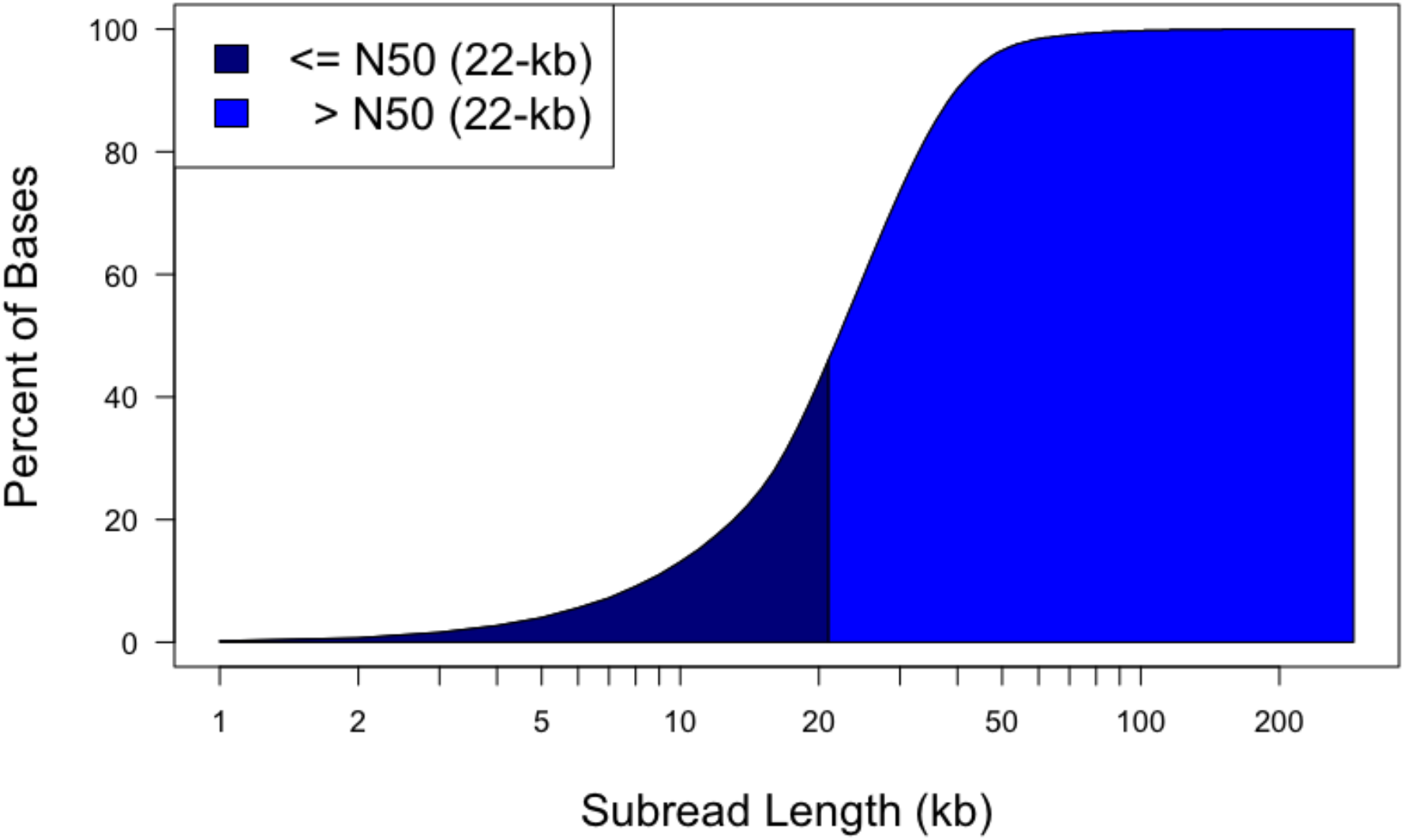
Cumulative distribution of subread lengths for Sequel II 8M SMRT Cell of 15kb size-selected library. Data were bioinformatically filtered prior to assembly to remove reads shorter than 500-bp and retain one subread per library molecule (see methods).

## Author contributions

SK performed assembly and curation. SG: performed genomic extraction and assembly curation. JU: performed sample collection. PB and CL performed library preparation and sequencing. JK, SK, SG and JU wrote the paper. SK, PB, AC, BC, BS, KH, JK, and SG conceived of and designed the project.

## Acknowledgments

We would like to thank Q.Wu (University of Science and Technology of China) and F. Cui (Institute of Zoology, Chinese Academy of Sciences) for sharing technical details about their previous genome assembly studies. We thank Angela Kauwe for assistance in the wetlab at USDA-ARS Hilo and Erica Smyers for providing the photograph for Figure 1B.

